# Competition-induced metabolic switching governs the interaction network in plant microbiome assembly

**DOI:** 10.64898/2026.01.26.701895

**Authors:** Hidehiro Ishizawa, Rikuya Umemoto, Nanoka Yoshida, Miki Furuya, Yosuke Tashiro, Daisuke Inoue, Michihiko Ike, Masahiro Takeo, Hiroyuki Futamata

## Abstract

Plants are colonized by characteristic microbiomes that help maintain plant health and function. However, the mechanisms that make these communities assemble in a consistent, repeatable manner remain poorly understood. Here, we show how microbial metabolic strategies for host-derived substrates drive plant microbiome assembly, using a six-member synthetic community that captures the taxonomic diversity of natural duckweed microbiome. Contrary to expectations of metabolic niche partitioning, five of six strains adopt similar metabolic states when colonizing the host alone, consistent with the preferential use of a shared substrate, acetaldehyde. In contrast, during competition with specific community members, these strains consistently switch toward alternative substrates, including sugars and aromatics. Strikingly, this metabolic switching accompanies all negative interspecies interactions observed in the community, indicating that a single class of metabolic response dominates the inhibitory interaction network organizing community structure. By comparison, although metabolite cross-feeding is widespread, its contribution to facilitative interactions depends on the background of resource competition. Together, these findings highlight competition-induced metabolic switching as a primary driver of microbial interaction networks, and provide a cross-scale account of how microbial metabolic strategies underpin the reproducible assembly of plant microbiomes.

## INTRODUCTION

Plant roots and leaves create favorable microenvironments that foster diverse microbial colonization, forming complex communities collectively referred to as the plant microbiome (Turner et al., 2013; Trivedi et al., 2020). Although microbiome structure is shaped by local environmental conditions rather than being predetermined, many plant hosts reproducibly assemble conserved core microbiomes that are stably maintained within host environments (Lundberg et al., 2012; Toju et al., 2018). These communities can enhance plant growth and stress tolerance (Ritpitakphong et al., 2016; Moore et al., 2023), whereas their disruption is frequently linked to disease susceptibility and nutrient imbalance (Ginnan et al., 2020; Lee et al., 2021; Mukai et al., 2024). Despite extensive community profiling documenting taxonomic consistency across plant microbiomes (Walters et al., 2018; Thiergart et al., 2019; Saimee et al., 2025), the mechanistic basis of how stable microbiome structures emerge and persist remains incompletely understood.

Metabolic interactions mediated by host-derived substrates—including resource competition, niche partitioning, and cross-feeding—are widely considered major drivers of microbiome assembly, in both plant and animal systems (Culp and Goodman, 2023; Medlock et al., 2018; Zhalnina et al., 2018; Sasse et al., 2018; Yang et al., 2025; Feng and Liu, 2025). However, meta-omics surveys typically provide static snapshots of resource use, often overlooking context-dependent metabolic switching that can alter individual fitness and interaction strengths. Consequently, mechanistic links between microbial metabolism and ecological outcomes, such as interspecies interactions and community structure, remain largely elusive. Recent efforts using genome-scale metabolic modeling (Schäfer et al., 2023) or gnotobiotic two-member systems (Hemmerle et al., 2022) have begun to address this gap, but fundamental questions persist: How is microbial metabolism remodeled in response to competitors? How does such metabolic plasticity scale up to ecological interaction networks that shape community structure? And to what extent do metabolic interactions contribute to assembly outcomes alongside non-metabolic mechanisms such as toxin-mediated antagonism and spatial competition?

Here, we address these questions using a six-member synthetic community (SynCom) associated with duckweed (*Lemna japonica*), an emerging model system for plant-microbe interactions (Acosta et al., 2021; Ishizawa et al., 2025). This SynCom captures the taxonomic diversity of the natural duckweed microbiome, with each strain representing one of six dominant bacterial families (Ishizawa et al., 2020a). Clonal propagation of duckweed stabilizes system-level physiology (Datko et al., 1980), thereby minimizing developmental confounding that complicates metabolic profiling in other plant systems (Xiong et al., 2021; Bourceret et al., 2022). Moreover, strain-specific fitness and interspecies interactions in this SynCom have been comprehensively quantified, allowing the prediction of community structure using a Lotka-Volterra-type model (Ishizawa et al., 2024a). Together, these features provide a tractable platform for integrating molecular and ecological measurements, thereby enabling a cross-scale mechanistic account of plant microbiome assembly.

We generated comprehensive condition-specific transcriptomes across all mono-inoculation, pairwise, and community contexts, complemented by exometabolomics and functional validation using knockout mutants. By integrating these molecular profiles with quantified interaction coefficients, we show that competition-driven metabolic switching—a strategy shared across phylogenetically diverse taxa—is the primary determinant of the inhibitory interaction network that gives rise to consistent community structure. Our findings provide a cross-scale, mechanistic view of how microbial metabolic interactions translate into ecological patterns in plant microbiomes.

## RESULTS

### Duckweed-based SynCom

The six-member SynCom established by Ishizawa et al. (2020) was used in this study. This SynCom represents a reproducible community structure on the duckweed surface, with *Acidovorax* sp. DW039, *Methylophilus* sp. DW102, and *Asticcacaulis* sp. DW145 as dominant strains, whereas *Chryseobacterium gambrini* DW100, *Herbaspirillum* sp. DW155, and *Novosphingbium olei* DW067 are present as minor members, reflecting the family-level structure of the natural duckweed microbiome (Fig. 1a). The colonization density of each species in the SynCom was largely consistent with the pairwise interspecies interaction network (Fig. 1b); species receiving negative interactions (e.g., DW039, DW067, and DW155) exhibited reduced abundance in the SynCom whereas those receiving positive interactions (e.g., DW100 and DW102) maintained robust abundance even in the community context (Ishizawa et al., 2024a). The sole exception was strain DW145 whose reduced colonization density in the SynCom was explained by higher-order interactions, whereby the co-presence of DW145, DW102, and another strain resulted in a pronounced decline of DW145 abundance. The resulting community structure was quantitatively captured using a Lotka–Volterra-type model calibrated with pairwise and higher-order interaction coefficients (Ishizawa et al., 2024a), suggesting that elucidating the mechanistic basis of these interactions can provide a molecular-to-ecosystem-scale account of plant microbiome assembly.

**Fig. 1.**
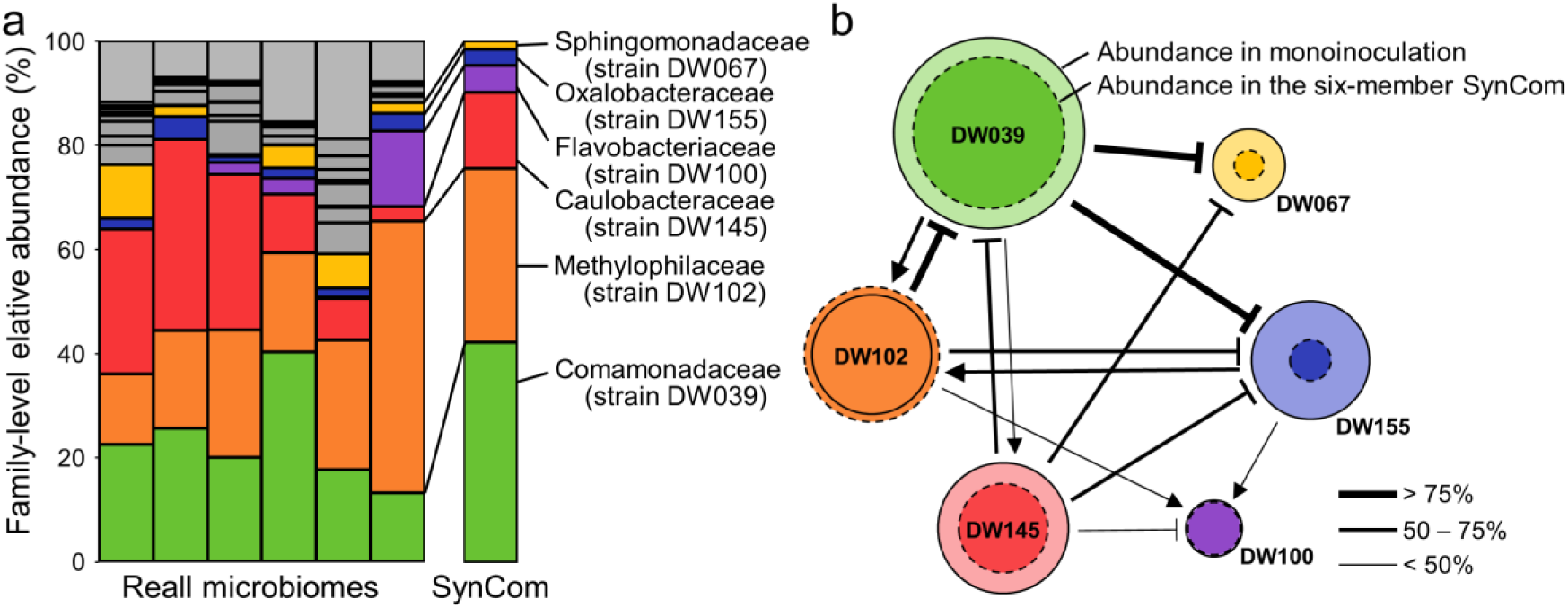
Properties of the duckweed-based synthetic community (SynCom) have been characterized previously. (a) Comparison of bacterial community structure between six duckweed microbiomes profiled using 16S rRNA gene amplicon sequencing (Ishizawa et al., 2020b) and the six-member SynCom profiled via colony counting (Ishizawa et al., 2024). Both studies used the same duckweed clone and culture conditions, but differed in inoculum source (natural freshwater microbiomes versus a mixture of isolates composing the SynCom). (b) Pairwise interactions among SynCom members based on Ishizawa et al. (2024a). Solid and dashed circles indicate colonization density (CFU per mg plant fresh weight) under mono-inoculation and in the six-member community, respectively. Arrows indicate significant pairwise interactions, defined as changes in colonization density under pairwise co-inoculation; pointed and blunt arrowheads represent positive and negative effects, respectively. Arrow thickness represents the percent change in colonization density relative to mono-inoculation.

### Ability of SynCom members to utilize host-derived substrates

To determine whether interspecies interactions can be explained by the differential utilization of host-derived substrates, we cultured the six strains separately in water-extracted duckweed exudates as the sole carbon source. After 48 h of cultivation, a duration deemed sufficient for the consumption of available substrates, culture supernatants were analyzed using untargeted GC-MS metabolomics.

GC-MS detected 1,081 valid metabolic peaks, of which 184 were putatively annotated based on retention time and mass spectrum similarity (Fig. 2a). Detected metabolites included primary metabolites, such as sugars, organic acids, and amino acids, as well as secondary metabolites, including aromatics, terpenoids, amines, and steroids (Supplementary Data S1). Among the 1,081 peaks, 196 were significantly depleted or enriched by at least one bacterial strain compared with uninoculated controls (ANOVA with Dunnett post-hoc test, *P* < 0.05), likely reflecting consumption and production during cultivation, respectively.

**Fig. 2.**
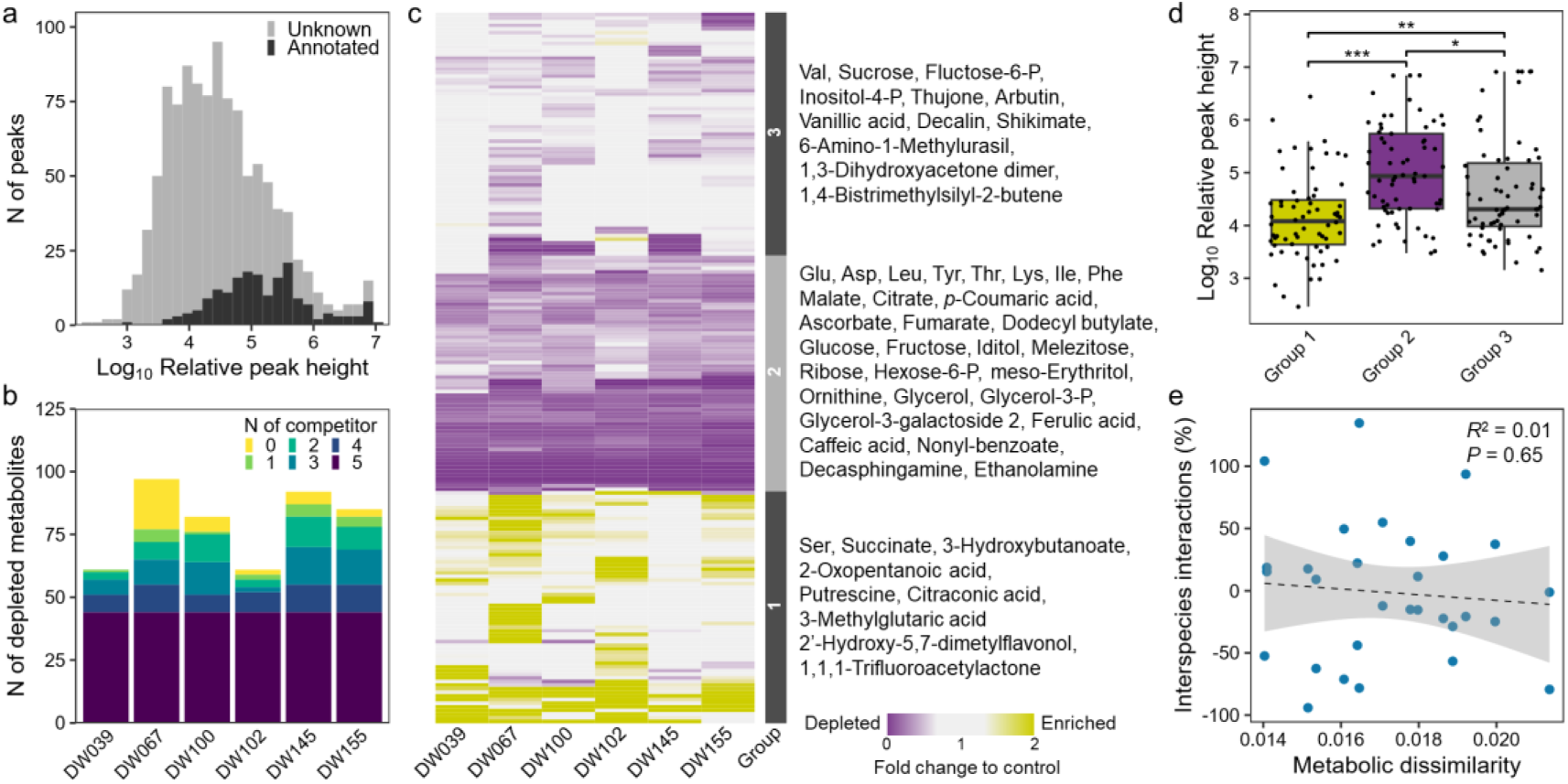
GC–MS analysis of culture residues from SynCom members grown in duckweed exudates. (a) Histogram of the estimated abundance (relative peak height) of 1,081 metabolic peaks in uninoculated control samples; 184 peaks were putatively annotated. (b) Number of depleted metabolic peaks per strain. Colors indicate the number of SynCom members depleting the same peak (0, depleted only by that species; 5, depleted by all SynCom members). (c) Heatmap of 196 metabolic peaks significantly depleted or enriched by at least one bacterial strain. Fold changes are shown relative to uninoculated control. Hierarchical clustering identified three groups: enriched metabolites (Group 1), metabolites depleted by most strains (Group 2), and metabolites depleted by a subset of species (Group 3). Annotated metabolites for each group are listed. (d) Relative peak heights of metabolic peaks classified into the three groups. Asterisks indicate significance: *, *P* < 0.05; **, *P* < 0.01, ***, *P* < 0.001. (e) Relationship between metabolic profile dissimilarity and pairwise interaction strength.

The range of consumable metabolites largely overlapped among strains, with most available metabolites serving as “public goods” consumed by multiple community members (Fig. 2b). These shared metabolites, classified as Group 2 via hierarchical clustering (Fig. 2c), included proteogenic amino acids, organic acids, sugars, glycerol derivatives, and lignin precursors (i.e., caffeic acid and ferulic acid). Although several Group 3 metabolites were consumed exclusively by subsets of strains, these metabolites generally exhibited lower relative peak heights than those in Group 2 (Fig. 2d). Together, these findings indicate that the six SynCom members are inherently in competitive relationships toward duckweed exudates despite their stable coexistence on the host.

Next, we tested whether metabolic similarity accounted for the strength of interspecies interactions. Intuitively, bacterial pairs with grater overlap in metabolic capacity would be expected to experience stronger competition, resulting in inhibitory interactions (Schäfer et al., 2023; Schlechter et al., 2023). However, similarity in metabolic profiles did not correlate with previously quantified interaction coefficients (Fig. 2e), indicating that static metabolic capacity toward host-derived substrates does not simply predict the ecological dynamics underlying community assembly.

### Transcriptome profiling under competitive and noncompetitive conditions

To investigate dynamic metabolic interactions among the six strains, we performed RNA-seq on 22 duckweed culture systems in triplicates: mono-inoculations of each strain, all 15 pairwise combinations, and the full six-member community. After 10 days of co-cultivation, total bacterial mRNA was extracted from duckweed surfaces and sequenced, and the reads were mapped to the reference genomes of the six strains. Sequencing depth varied largely due to differences in colonization density on the duckweed surface; however, sufficient mapped reads were obtained to identify major metabolic pathways (approximately 0.3 to 47 million reads per strain), except from a single replication of the full SynCom system, which yielded only < 0.13 million reads for strains DW067, DW100, and DW145 (Supplementary Data S2). Consequently, only transcriptomic profiles with >0.3 million reads were included in subsequent analyses, resulting in replication numbers of *n* = 3 (39 profiles) or *n* = 2 (3 profiles).

Standard differential expression analysis identified 57,872 differentially expressed gene calls (FDR < 0.05, fold change > 2), corresponding to 6,881 unique genes across conditions (Supplementary Data S3), which precluded systematic gene-by-gene interpretation. We therefore visualized expression patterns using self-organizing map (SOM) clustering, which groups genes based on co-expression across conditions for each strain (Figs. 3–5). For clarity, SOM analyses focused on metabolism-related genes from KEGG metabolic pathways (Kanehisa et al., 2000), while non-metabolic genes were analyzed separately (Fig. S1). Gene-to-cluster assignments are provided in Supplementary Data S4. Functional enrichment analysis using Fisher’s exact test identified 560 significantly enriched KEGG categories (*P* < 0.05) across SOM clusters (Supplementary Data S5).

**Fig. 3.**
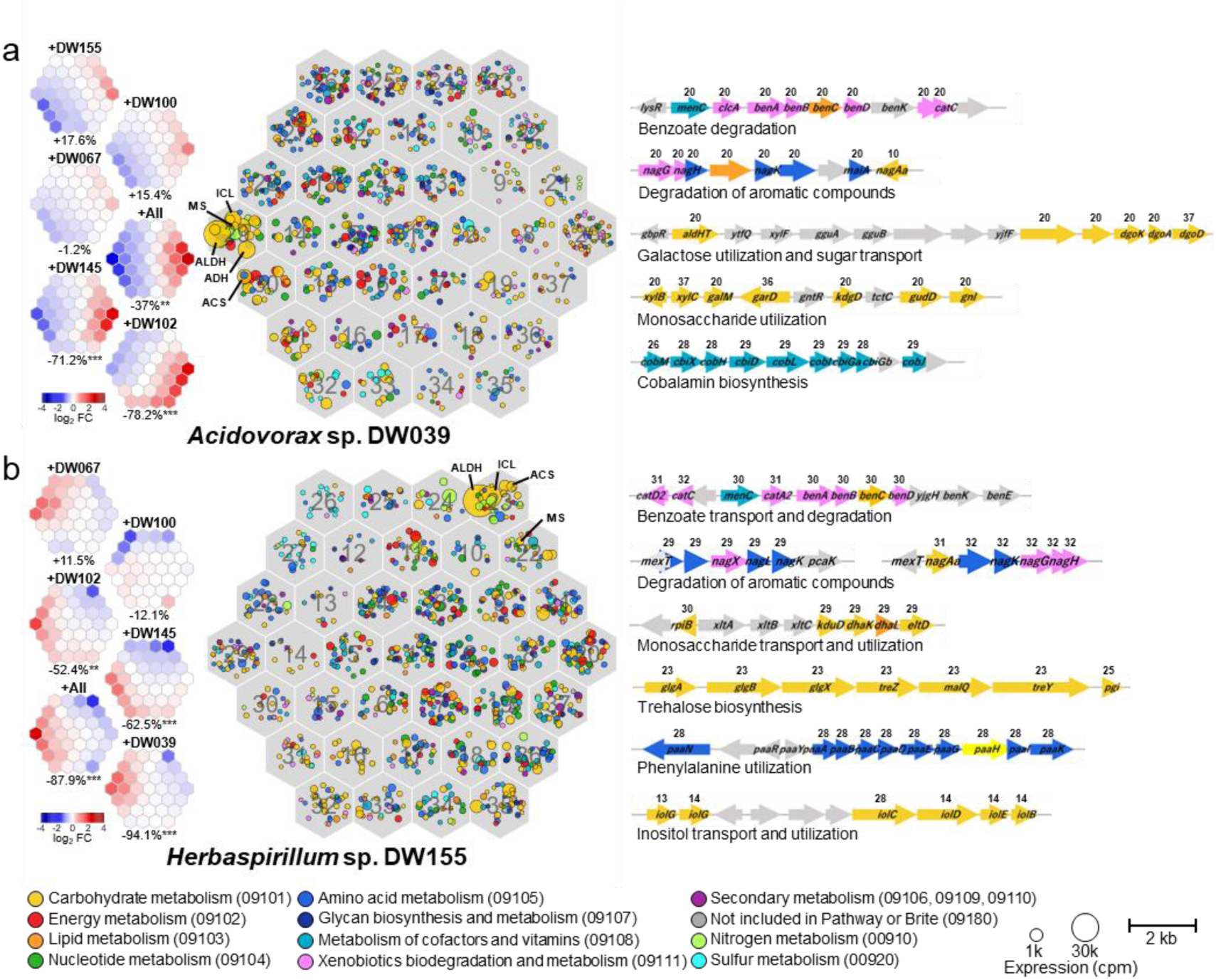
Metabolic responses of (a) *Acidovorax* sp. DW039 and (b) *Herbaspirillum* sp. DW155 across community contexts (pairwise co-inoculation and the six-member SynCom). Central hexagonal maps show self-organizing map (SOM) clustering based on gene expression patterns. Each small hexagon represents one SOM cluster; symbols denote genes, with colors indicating KEGG functional categories and sizes reflecting expression levels under mono-inoculation. Heatmaps show mean gene expression levels (fold change relative to mono-inoculation) within each SOM cluster. Values below the heatmaps indicate percent changes in colonization density relative to mono-inoculation. Right panels highlight representative gene clusters with altered expression under co-inoculation; numbers above genes denote SOM cluster assignments. ALDH, aldehyde dehydrogenase; ACS, acetyl-CoA synthetase; ICL, isocitrate lyase; MS, malate synthase.

**Fig. 4.**
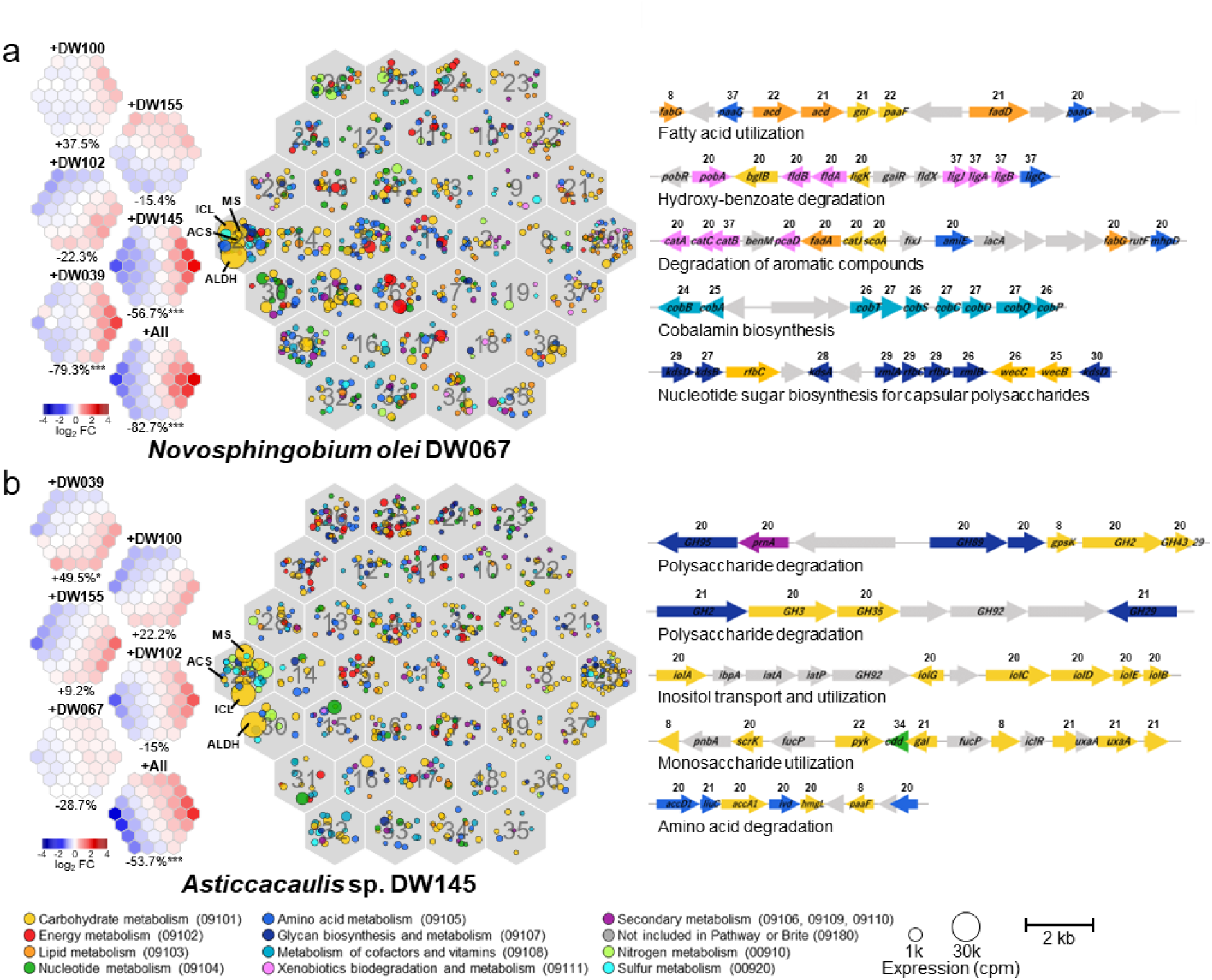
Metabolic responses of (a) *Novosphingobium olei* DW067 and (b) *Asticcacaulis* sp. DW145 across community contexts (pairwise co-inoculation and the six-member SynCom). Central hexagonal maps show self-organizing map (SOM) clustering based on gene expression patterns. Each small hexagon represents one SOM cluster; symbols denote genes, with colors indicating KEGG functional categories and sizes reflecting expression levels under mono-inoculation. Heatmaps show mean gene expression levels (fold change relative to mono-inoculation) within each SOM cluster. Values below the heatmaps indicate percent changes in colonization density relative to mono-inoculation. Right panels highlight representative gene clusters with altered expression under co-inoculation; numbers above genes denote SOM cluster assignments. ALDH, aldehyde dehydrogenase; ACS, acetyl-CoA synthetase; ICL, isocitrate lyase; MS, malate synthase.

**Fig. 5.**
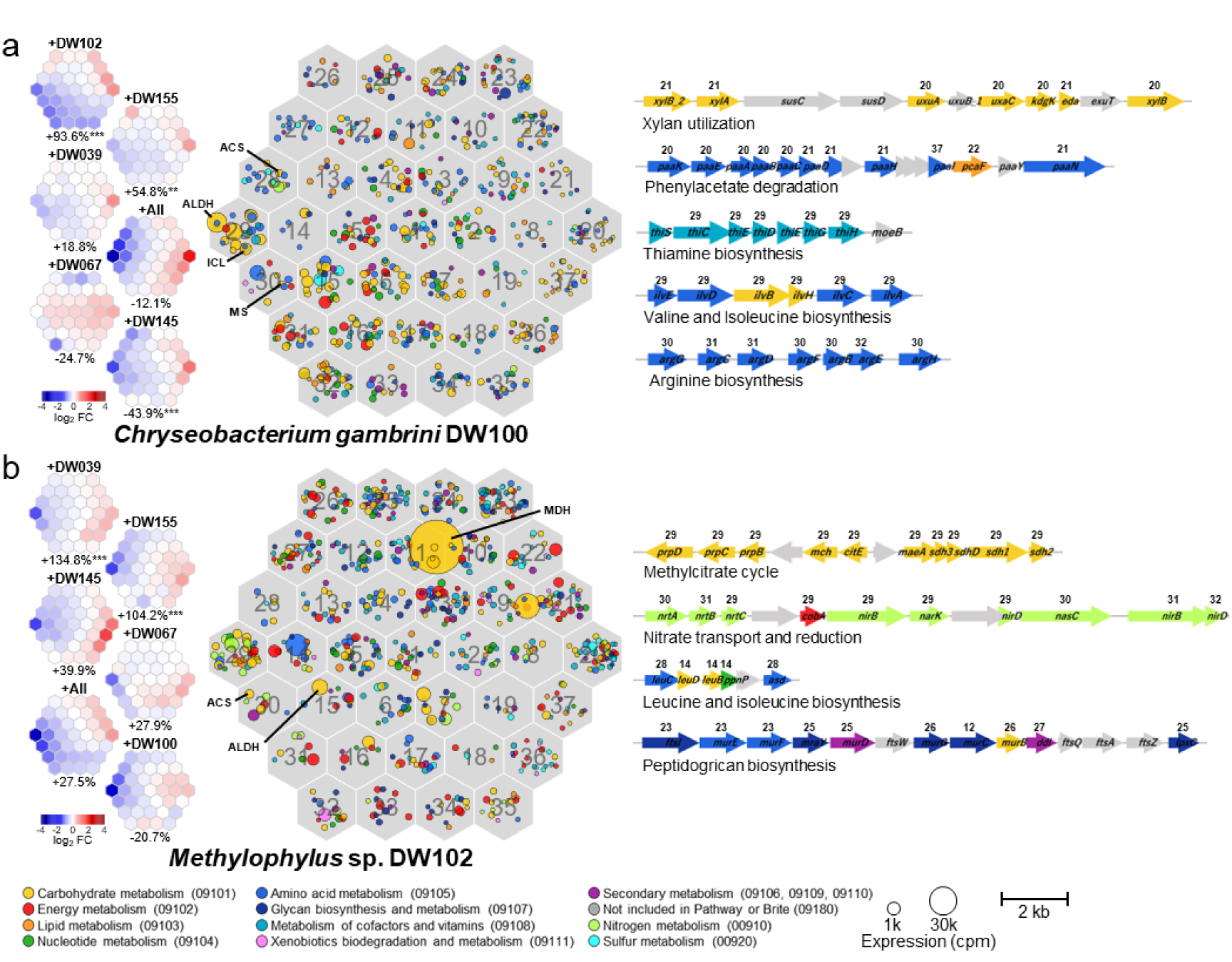
Metabolic responses of (a) *Chryseobacterium gambrini* DW100 and (b) *Methylophilus* sp. DW102 across community contexts (pairwise co-inoculation and the six-member SynCom). Central hexagonal maps show self-organizing map (SOM) clustering based on gene expression patterns. Each small hexagon represents one SOM cluster; symbols denote genes, with colors indicating KEGG functional categories and sizes reflecting expression levels under mono-inoculation. Heatmaps show mean gene expression levels (fold change relative to mono-inoculation) within each SOM cluster. Values below the heatmaps indicate percent changes in colonization density relative to mono-inoculation. Right panels highlight representative gene clusters with altered expression under co-inoculation; numbers above genes denote SOM cluster assignments. ALDH, aldehyde dehydrogenase; ACS, acetyl-CoA synthetase; ICL, isocitrate lyase; MS, malate synthase.

### Metabolic strategies of *Acidovorax* sp. DW039 and *Herbaspirillum* sp. DW155

Transcriptomic profiling revealed a shared metabolic response among the SynCom members, including the two Burkholderiales strains DW039 and DW155 (Fig. 3). Under competition, these strains downregulated key genes involved in central carbon metabolism, including aldehyde dehydrogenase (ALDH), acetyl-CoA synthetase (ACS), isocitrate lyase (ICL), and malate synthase (MS). ALDH was the most highly expressed metabolic gene under mono-inoculation but decreased by up to ca. 330-fold in strain DW039 and ca. 7.5-fold in strain DW155 in the presence of specific competitors. This downregulation was most pronounced when colonization density was strongly suppressed (Fig. 3), indicating competition-driven metabolic switching away from the primary pathway.

Concomitant with the down regulation of ALDH, ACS, ICL, and MS, alternative metabolic pathways were upregulated under competition. In strain DW039, several gene clusters associated with aromatic degradation and monosaccharide utilization were upregulated under competition (Fig. 3a). The most strongly upregulated SOM cluster (cluster 20) was enriched for KEGG categories including “Benzoate degradation,” “Pentose and glucuronate interconversions,” “Xylene degradation,” “Fatty acid degradation” (Supplementary Data S3), consistent with switching toward the utilization of aromatics, monosaccharides, and fatty acids. Strain DW155 showed a similar upregulation of genes involved in aromatic and monosaccharide utilization under competition (Fig. 3b). Together, these observations suggest that both strains strategically alter their carbon source preferences in response to strong competitors, enabling stable coexistence albeit at reduced abundance. The pronounced similarity in their metabolic strategies may underlie the particularly strong inhibitory effect of strain DW039 on strain DW155.

Both strains also exhibited transcriptional signatures indicative of metabolic cross-feeding. In strain DW039, biosynthetic genes for cobalamin and some amino acids (Arginine, tryptophan, and lysine) were downregulated in the presence of other bacteria. In parallel, both strains downregulated nitrate transporters and nitrate reductases while upregulating transporters for amino acids, urea, and polyamines during co-inoculation (Fig. S1), indicating a shift in nitrogen source from nitrate present in the culture medium to nitrogenous metabolites likely supplied by other bacteria. Additionally, both strains increased expression of genes involved in galactose, xylose, and fucose utilization when co-inoculated with strains DW100 and DW145. Because strains DW100 and DW145 harbor multiple genes encoding plant polysaccharide degrading enzymes (Ishizawa et al., 2024b), these expression patterns are consistent with cross-feeding interactions wherein strains DW100 and DW145 depolymerize plant-derived polysaccharides, and strains DW039 and DW155 subsequently utilize the released sugars.

### Metabolic strategies of *Novosphingobium olei* DW067 and *Asticcacaulis* sp. DW145

The downregulation of ALDH, ACS, ICL, and MS was consistently observed in the two Alphaproteobacteria strains, DW067 and DW145 (Fig. 4). In DW067, strong ALDH downregulation (up to 71-fold) occurred specifically under conditions of reduced colonization density and was accompanied by the upregulation of two operons involved in aromatic hydrocarbon degradation (Fig. 4a; Supplementary data S4). The most strongly upregulated SOM cluster (cluster 20) was enriched for genes in “Fatty acid degradation” and “Valine, leucine and isoleucine degradation” KEGG categories (Supplementary Data S3), consistent with switching to aromatics, fatty acids, and amino acids under competition. In contrast, DW145 exhibited only weak responses to pairwise competition, with limited ALDH downregulation (<3.4-fold) and modest reduction in colonization density (≤ 28.7%). However, strong responses were observed in the six-member community, including 19-fold ALDH downregulation and 53.7% decrease in colonization density (Fig. 4b). This pattern indicates the dominance of higher-order interactions involving ≥3 species rather than simple pairwise effects (Ishizawa et al., 2024a). Under competitive conditions, DW145 upregulated genes encoding glycoside hydrolases and monosaccharide utilization pathways, consistent with switching to plant-derived polysaccharides such as mucilage and pectin.

Both strains also exhibited transcriptional signatures consistent with metabolic cross-feeding. During co-inoculation, genes involved in sulfate and nitrate assimilation were downregulated in both strains, suggesting reliance on organic sulfur and nitrogen compounds released by other community members. In strain DW067, cobalamin biosynthesis genes were downregulated, except when paired with DW155. This strain also exhibited pronounced downregulation of genes associated with motility and capsular polysaccharide synthesis when colonization density declined, consistent with the importance of these traits for bacterial colonization of duckweed surfaces (Ishizawa et al., 2022).

### Metabolic strategies of *Chryseobacterium gambrini* DW100 and *Methylophilus* sp. DW102

The Flavobacteriaceae strain DW100 also downregulated ALDH, ACS, ICL, and MS under competitive conditions, yet maintained relatively high colonization density except when paired with strain DW145 (Fig. 5a). This robustness likely reflects the comparatively low basal expression of these genes under mono-inoculation. Instead, strain DW100 constitutively expressed genes involved in polysaccharide degradation at high levels, aligning with the reported role of *Flavobacteriaceae* in degrading pectin and hemicellulose in plant microbiomes (McBride et al., 2009; Kolton et al., 2013). The decreased colonization density observed during co-inoculation with strain DW145 likely results from the combined effects of ALDH downregulation and metabolic niche overlap, as DW145 also utilizes plant-derived polysaccharides.

The methylotrophic Gammaproteobacteria strain DW102 (*Methylophilus* sp.) expressed exceptionally high levels of methanol dehydrogenase (Fig. 5b), indicating a strong dependence on methanol released during plant pectin degradation (Sy et al., 2005; Dorokhov et al., 2018; Yurimoto and Sakai, 2023). This metabolic specialization likely explains why strain DW102 was the only strain not inhibited by other community members, as it monopolizes methanol as a primary carbon source and thereby avoids competition for more widely utilized substrates. Under co-inoculation, strain DW102 strongly downregulated genes involved in methylcitrate cycle, which mediates the assimilation and detoxification of propionyl-CoA generated from specific carbon sources such as branched-chain amino acids and odd-chain fatty acids (Dolan et al., 2018; Huang et al., 2023). This expression pattern is consistent with the reduced use of these supplementary carbon sources in the presence of other heterotrophs.

Both strains exhibited signatures of cross-feeding interactions. Under competitive conditions, strains DW100 and DW102 downregulated biosynthetic pathways for thiamine and several amino acids (Fig. 5), indicating increased dependence on other community members for these metabolites. Conversely, DW102 upregulated genes involved in folate biosynthesis (cluster 20; Fig. 5b; Supplementary Data S4), suggesting self-sufficiency for this vitamin. Strain DW102 also exhibited partner-specific regulation of genes associated with motility and peptidoglycan biosynthesis (Fig. S1). Notably, several uncharacterized genes in DW100 were upregulated during facilitation by DW102, indicating that non-metabolic mechanisms may also contribute to this positive interaction.

### Role of the ALDH gene in the duckweed environment

Transcriptomic analyses showed that five of the six SynCom members (excluding DW102) strongly expressed the ALDH, ACS, ICL, and MS genes during mono-inoculation, but markedly downregulated these genes when colonization density decreased due to inhibitory interactions (Fig. 6a). In these five strains, ALDH genes (commonly annotated as KEGG K00138) were the most highly expressed metabolic genes, indicating a central role in duckweed colonization. Comparative sequence analysis showed that ALDH genes shared more than 62.9% amino acid identity and were frequently co-localized with genes encoding alcohol dehydrogenase and DUF779 domain proteins (Fig. 6b). Additionally, genes encoding a Fis family transcriptional regulator were present within the ALDH gene clusters of four strains (DW039, DW067, DW145, and DW155), all of which exhibited particularly strong ALDH downregulation under competitive conditions.

**Fig. 6.**
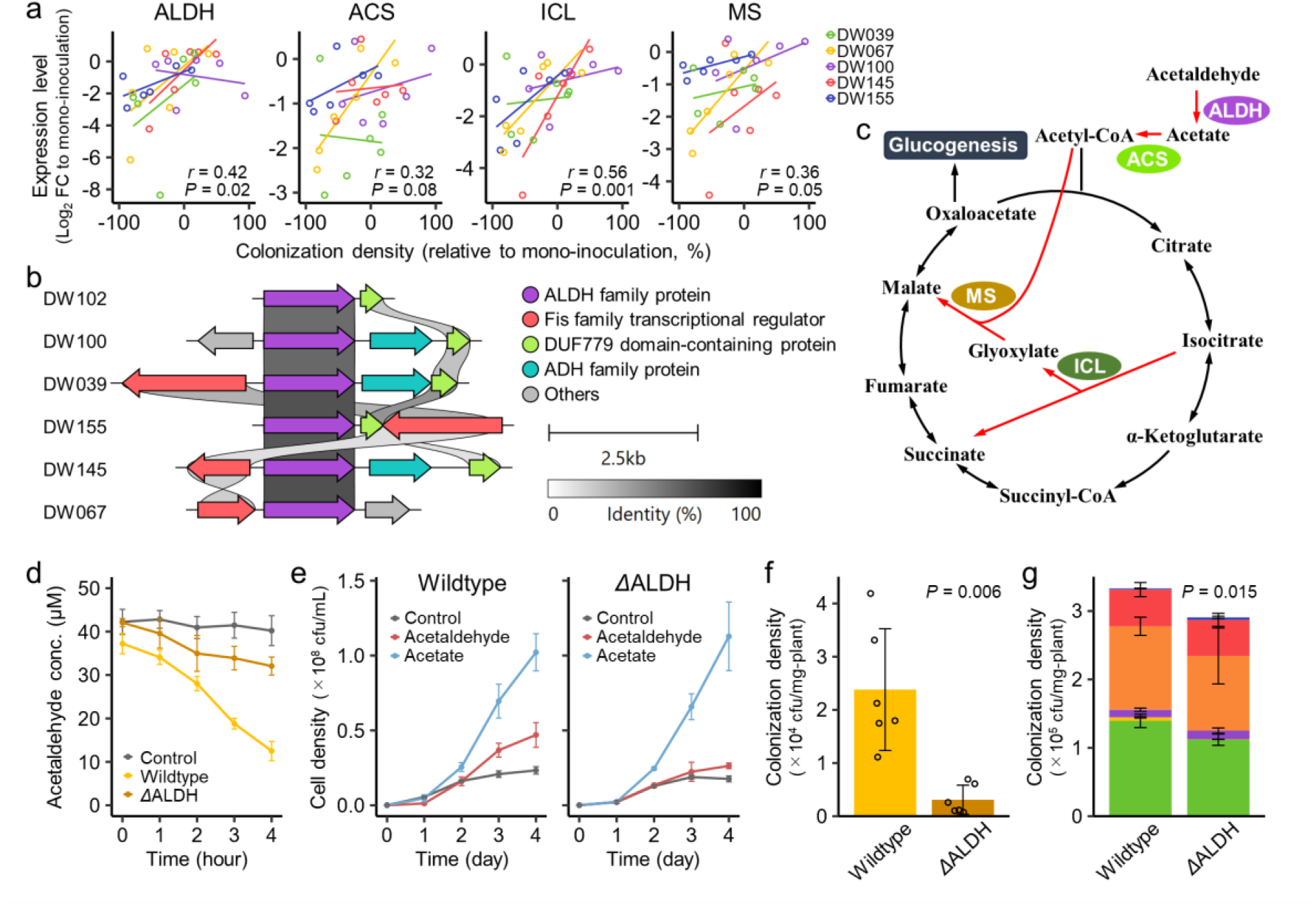
Functional role of aldehyde dehydrogenase (ALDH) in duckweed colonization. (a) Expression levels of ALDH, acetyl-CoA synthetase (ACS), isocitrate lyase (ICL), and malate synthase (MS) positively correlate with colonization density across five SynCom members. (b) Synteny of the ALDH gene cluster among SynCom members; ribbons indicate amino acid sequence similarity. (c) Proposed metabolic pathway linking acetaldehyde utilization to central carbon metabolism via ALDH, ACS, ICL, and MS. (d) Acetaldehyde utilization by *Novosphingobium olei* DW067 wild-type and ΔALDH mutant in liquid culture. (e) Growth curves in minimal medium supplemented with 0.01% yeast extract and either 50 µM acetaldehyde or 50 µM acetate. (f) Colonization density of DW067 wild-type and ΔALDH mutant after 10 days of mono-inoculation on gnotobiotic duckweed. (g) Community structure of the six-member SynCom containing eigher strain DW067 wild-type or ΔALDH mutant. Colors correspond to Fig. 1a. Error bars indicate standard deviations.

ALDH and ACS catalyze the sequential conversion of acetaldehyde to acetate and acetyl-CoA, respectively, whereas ICL and MS constitute the glyoxylate shunt, which enables assimilation of C2 compounds by bypassing the two decarboxylation steps in the TCA cycle (Fig. 6c) (Kornberg and Madsen, 1958). Therefore, we hypothesized that the high expression of these genes reflects preferential utilization of acetaldehyde—a common C2 aldehyde—on the duckweed surface. To test this hypothesis, an ALDH knockout mutant of strain DW067 was constructed and its phenotype was compared with the wild-type. Consistent with the expected role of ALDH in converting acetaldehyde to acetate, the mutant exhibited impaired consumption and growth on acetaldehyde while maintaining normal growth on acetate *in vitro* (Fig. 6de). The mutant showed significantly reduced colonization of duckweed under both mono-inoculation and the full SynCom conditions, whereas ALDH deletion did not substantially affect the colonization densities of other SynCom members (Fig. 6f). Acetaldehyde was detected in the culture medium of aseptically cultivated duckweed at 0.12 ± 0.06 µM. Although this concentration is low, continuous acetaldehyde release from plant surface likely provides a sustained substrate supply for bacterial growth. Together, these results support the hypothesis that ALDH is essential for efficient acetaldehyde utilization and for achieving high population densities on the duckweed surface.

To assess whether ALDH contributes to the utilization of other duckweed-derived compounds, we compared exometabolomic profiles of the strain DW067 wild-type and ΔALDH mutant grown in extracted duckweed exudates. The resulting profiles were highly similar between strains (Fig. S2; Supplementary Data S5), and no metabolite peaks were significantly more depleted in the wild-type than in the mutant (one-way ANOVA, FDR <0.05). Acetaldehyde itself was not detected, likely due to evaporation during sample lyophilization.

## DISCUSSION

Metabolic interactions among sympatric microbes, including resource competition, niche partitioning, and cross-feeding, are widely recognized as key drivers of microbiome assembly (Baran et al., 2015; Goldford et al., 2018; Blasche et al., 2021). However, how these interactions emerge and drive ecological processes underlying community assembly remains poorly understood, limiting our ability to predict and control microbiome dynamics. By directly linking species-resolved metabolic strategies with quantified ecological parameters in a duckweed SynCom, we show that five of six members, representing core duckweed-associated families, exhibit similar metabolic states under mono-inoculation, likely reflecting preferential utilization of the host-derived acetaldehyde. In contrast, when coexistence with specific competitors, these strains consistently undergo metabolic switching toward alternative substrates, including sugars, lipids, and aromatics. Across the interspecies interaction network observed in this SynCom, such competition-induced metabolic switching consistently accompanies negative effects, suggesting that a broadly shared metabolic response to resource competition organizes the inhibitory network underlying reproducible community structure.

Inhibitory interactions among the SynCom members were always unidirectional, with no mutual inhibition or rock–paper–scissors cycles often associated with ecosystem stability (Kerr et al., 2002) (Fig. 1a). Furthermore, under pairwise competition, metabolic switching characterized by pronounced ALDH downregulation was typically observed in only one of the competing species (Figs. 3–5), suggesting a clear competitive hierarchy for shared metabolic niches (Higgins et al., 2017). Although microbial community assembly is often treated as stochastic, with species arrival order and ecological drift generating divergent outcomes (Debray et al., 2022; Hayashi et al., 2024), the highly consistent community structures reported across plant microbiomes remain difficult to explain. While classical niche theory emphasizes innate niche differentiation as a basis for community stability, our findings indicate that most strains initially prefer similar metabolic niches. Instead, hierarchical competitive strength, coupled with realized niche divergence driven by competition-induced metabolic switching, underpins stable community structure. Consistent with this interpretation, Hemmerle et al. (2022) reported competition-induced metabolic reprogramming in pairwise *Sphingobium*-*Rhizobium* interactions in the gnotobiotic *Arabidopsis* phyllosphere. Together, these observations suggest that competition-driven metabolic switching may be widespread across plant–microbiome systems, providing a mechanistic basis for consistent taxonomic structures.

Cross-feeding is commonly regarded as a key driver of facilitative interactions in microbial communities (Oña et al., 2021; Culp and Goodman, 2023). Although transcriptome profiles revealed numerous signatures consistent with metabolite cross-feeding, particularly involving cofactors, nitrogen sources, and polysaccharide degradation (Figs. 3–5), these signatures were broadly distributed across many strain combinations and did not specifically predict the facilitative interactions identified in this system. This pattern suggests that the benefits of cross-feeding are strongly context-dependent, becoming apparent only when resource competition is sufficiently relaxed. Additionally, specific strain combinations induced changes in motility and biofilm formation (Fig. S1), suggesting that shifts in bacterial life-history strategies may also contribute to facilitation. Indirect mechanisms, such as the modulation of plant immunity (Teixeira et al., 2021), could provide alternative explanations.

Although pairwise interactions provide a foundation for understanding community assembly (Ortiz et al., 2021; Zhu et al., 2025), emergent phenomena not predictable from pairwise interactions alone, often referred to as higher-order interactions, may also play important roles (Sanchez et al., 2019; Sundarraman et al., 2020; Gibbs et al., 2022). In our duckweed SynCom, for example, strain DW102 had a nearly neutral effect on strain DW145 in pairwise competition but became strongly inhibitory in the presence of third-party species (Ishizawa et al., 2024a). At the transcriptomic level, strain DW102 altered the metabolic state of DW145 under pairwise competition, yet pronounced ALDH downregulation occurred only in the six-member SynCom (Fig. 4b), suggesting that higher-order effects on ALDH expression underlie emergent DW145 population dynamics. Additionally, most negative interactions in this SynCom act subadditively on species abundances, exerting weaker combined effects than predicted by additive models (Ishizawa et al., 2024a). This deviation can be explained by the shared mechanistic basis of inhibitory interactions, namely competition-induced metabolic switching, such that once a strain has switched its metabolic state, additional competitors induce only marginal further inhibition.

Our findings further identify acetaldehyde as a key host-derived substrate influencing metabolic strategies and population success among various duckweed-associated bacteria. Acetaldehyde is a common plant metabolic intermediate and, because of high membrane permeability, appreciable amounts may diffuse from plant tissues into surrounding environments (Jardine and McDowell, 2023). Its production increases under hypoxic conditions (Kimmerer and MacDonald, 1987), suggesting potential relevance in aquatic environments, deep root zones, and during flooding (Kreuzwieser et al., 2004; Rottenberger et al., 2008). However, because of its high volatility, toxicity, and resulting detection difficulties, acetaldehyde has received limited attention in studies of plant–microbe interactions. Notably, Padaki et al. (2025) reported that acetaldehyde is the major volatile organic compound released by the marine microalga *Phaeodactylum tricornutum* and is consumed by associated bacterial strains during co-culture. Future studies should therefore examine whether acetaldehyde-mediated interactions are widespread across both aquatic and terrestrial plant systems.

In summary, by integrating species-resolved metabolic measurements with quantified interaction parameters in a taxonomically representative duckweed SynCom, we connect condition-dependent metabolic states to ecological interactions and community-level outcomes. Phylogenetically diverse strains converge on shared acetaldehyde utilization in isolation but consistently switch to alternative substrates under competition, and this conserved switching response accounts for most inhibitory interactions organizing community structure in this system. In contrast, transcriptomic signatures consistent with cross-feeding were widespread but did not reliably predict facilitative effects, highlighting the context dependence of facilitation and the potential contribution of non-metabolic mechanisms. Although our system represents a simplified model, it demonstrates how metabolic information can be mapped onto interaction networks to explain key ecological processes underlying community assembly. Together with recent works showing that specific host-derived metabolites can steer plant microbiome structure and function (Pascale et al., 2020; Wen et al., 2023), our results further motivate targeting key substrates as a strategy for rational microbiome design.

## MATERIALS AND METHODS

### Culture conditions of plant and bacterial strains

Duckweed *Lemna japonica* (RDSC5512; formerly identified as *L. minor*) was used as the host plant. Plants were maintained under sterile conditions in modified Hoagland medium (MH medium) (Toyama et al., 2006).

Six bacterial strains comprising the SynCom described by Ishizawa et al. (2020a), *Acidovorax* sp. DW039, *Novosphingobium olei* DW067, *Chryseobacterium gambrini* DW100, *Methylophilus* sp. DW102, *Asticcacaulis sp*. DW145, and *Herbaspirillum* sp. DW155, were used in this study. For experiments, these strains were cultured in 10 mL R2A medium (supplemented with 2% methanol for DW102) at 25°C with shaking at 150 rpm, then harvested by centrifugation (10,000 × *g*, 3 min) and washed twice with sterile MH medium.

### Exometabolomics

Duckweed exudates were prepared by soaking 180–220 sterile duckweed fronds in 4 mL ultra-pure water in a 5 mL tube, followed by shaking at 200 rpm for 3 h in an ice bath. Extracts were filtered through a 0.45 µm filter and stored at –80°C until use. Prior to cultivation, extracts were lyophilized (FDS-1000; Eyela, Tokyo, Japan), dissolved in 1 mL half-strength MH medium, pooled into a single tube, and sterilized using a 0.2 µm filter. The total organic carbon of the resulting medium was 44.0 mg-C L^−1^, corresponding to an adequate carbon to nitrogen ratio (ca. 17.6), as measured using TOC-V_CSN_ (Shimadzu, Kyoto, Japan). The blank medium was also prepared using the same procedure without duckweed plants.

Each bacterial strain was inoculated to the wells of a 48-well plate containing 300 µL of the dissolved duckweed exudate at an initial OD_600_ of 0.001 and incubated at 28°C for 48 h with rotary shaking at 150 rpm (*n* = 5). Control samples (exudate medium without bacterial inoculation) and Blank samples (blank medium without bacterial inoculation) were prepared concurrently. Following cultivation, 100 µL of 0.02 g L^−1^ ribitol was added to each well as an internal standard. Samples were filtered through a 0.2 µm filter and stored at –80°C until analysis. Quality control (QC) samples were prepared by pooling 100 µL from each sample.

Trimethylsilyl derivatization was performed by dissolving lyophilized samples in 25 µL methoxyamine solution (20 mg mL^−1^ in pyridine), vortexed for 10 s, and incubated at 30°C for 90 min with shaking at 150 rpm. Subsequently, 25 µL N-Methyl-N-trimethylsilyltrifluoroacetamide (Sigma-Aldrich, St. Louis, MO, USA) was added, followed by vortexin for 10 s and incubation at 37°C for 30 min with shaking at 150 rpm.

The GC-MS analysis of all samples was performed using a GC-MS-AP-2020NX system (Shimadzu). Each analytical run began with an n-alkane mixed standard (pyridine, 2:1), followed by Blank and QC samples, after which experimental samples, including Control, were analyzed in a randomized order. QC samples were measured every six injections. GC-MS operating conditions are detailed in Table S1.

The alignment and metabolite identification of detected peaks were performed using MS-DIAL version 4.9.221218 (Tsugawa et al., 2015) with parameters summarized in Table S2. Metabolites were annotated based on mass spectral similarity and Kovats retention index against the GCMS DB-Public-KovatsRI-VS3 database, using a total score threshold of >80%. When multiple peaks were assigned to the same compound, the peak with the highest total score was retained. The peaks of inorganic molecules (e.g., phosphate) and internal standards (ribitol) were removed. Peak heights were standardized using QC samples to correct for sensitivity drift. Outliers arising from experimental procedures, peak detection, or alignment were identified using Grubbs’ test (*P* < 0.05), with at most one outlier removed per set of five replicates. Peaks whose mean height in any experimental system was less than four times that in the Blank sample were regarded as contamination and discarded. Finally, one-way ANOVA followed by Dunnett’s post hoc test (*P* < 0.05) was used to identify metabolite peaks whose heights differed significantly from those of the Control. The comparison of wild-type and mutant strain was also performed by the same procedures but using Welch’s t-test for comparison. Statistical analyses were performed in R version 4.1.0 using the “outlier” and “multcomp” packages.

### Transcriptome analysis of duckweed-attached bacteria

To obtain samples for transcriptome analysis, we constructed duckweed-based SynComs comprising one, two, or six bacterial strains, following exactly the procedure described by Ishizawa et al. (2024a). Briefly, bacterial cells were mixed at equal ratios and inoculated into sterile duckweed cultures in 100 mL flasks containing 60 mL MH medium at an initial OD_600_ of 0.0005. Co-cultures were incubated in a growth chamber for five days, after which 10 fronds were transferred to fresh MH medium and cultivated for another five days. For RNA extraction, all duckweed plants in each flask (approximately 40–50 fronds) were harvested, gently wiped with sterile paper towels, and immediately soaked in 1.5 mL RNAprotect Bacteria Reagent (Qiagen, Hilden, Germany), followed by vortexing for 1 min. Then, bacterial cells in supernatant were collected via centrifugation at 5,000 × *g* for 10 min. Total RNA was then extracted by resuspending the pellets in 1 mg mL^-1^ lysozyme and 30 mAU mL^-1^ of Proteinase K (Qiagen) in 100 µL TE buffer (30 mM Tris-HCl, 1 mM EDTA, pH = 8.0) and incubating for 10 min. The extracts were purified using the RNeasy Mini Kit (Qiagen) according to the manufacturer’s protocol.

The sequencing library was prepared using the NEBNext Ultra II Directional RNA Library Prep Kit for Illumina (New England Biolabs, Ipswich, MA, USA), and 100 bp single-end sequencing was performed on an Illumina Novaseq 6000 system (Illumina, San Diego, CA, USA). The obtained sequencing reads were mapped to the genome sequences of the six SynCom members (Ishizawa et al., 2024b) using Salmon version 1.6.0 (Patro et al., 2017). Read counts were normalized with edgeR version 4.8.0 (Robinson et al., 2010, and differentially expressed genes (satisfying FDR < 0.05 and fold change > 2) were identified using “glmQLFTest” function.

For visualization, genes were clustered based on their expression patterns, defined as log_2_ fold changes relative to the mono-inoculation across the six competitive conditions, using SOM implemented in the “supraHex” R package (Fang and Gough, 2014). Mean log_2_ fold changes were then calculated for genes within each SOM cluster and used to generate a heatmap. Functional enrichment within each SOM cluster was assessed using Fisher’s exact test to determine whether genes assigned to specific KEGG categories were significantly enriched in each cluster.

### Gene knockout

An ALDH gene mutant of strain DW067 was generated via double-crossover homologous recombination. First, approximately 1 kb regions upstream and downstream sequences of the ALDH gene were cloned into the suicide vector pK18mobsacB (Schäfer et al., 1994) using In-Fusion HD Cloning Kit (Takara Bio, Shiga, Japan). The resulting constructs were introduced into the wild-type strain DW067 via triparental mating with donor (*E. coli* HST08) and helper (*E. coli* DH5α/pRK2013) strains. Transformants were selected on R2A agar plates supplemented with streptomycin (50 µg mL^−1^) and kanamycin (20 µg mL^−1^). The second crossover was performed by plating a log culture of the transformants on a R2A agar containing 5% sucrose. The successful deletion of the ALDH gene was confirmed via PCR amplification and sequencing of the target regions. Primers and bacterial strains used in these procedures are summarized in Table S3.

### Characterization of the knockout strain

The ability of the strain DW067ΔALDH mutant to utilize acetaldehyde was evaluated by inoculating overnight cultures of mutant and wild-type strains into 10 mL sterile MH medium in vials at an initial OD₆₀₀ of 0.2. Acetaldehyde was added to a final concentration of 50 µM, and cultures were incubated for 4 h at 22°C with shaking at 100 rpm. To monitor the acetaldehyde concentration, 50 µL culture supernatant was sampled at 0, 1, 2, 3, and 4 h and stored at –20°C until analysis. Acetaldehyde concentrations were determined using an Acetaldehyde Assay Kit (Sigma-Aldrich) following removal of cells via centrifugation (10,000 × *g*, 5 min). Each condition included four biological replicates, including a non-inoculated control.

To assess the growth of strain DW067ΔALDH in the presence of acetaldehyde and its downstream metabolite acetate, mutant and wild-type strains were inoculated into 10 mL MH medium supplemented with 0.01% yeast extract and 50 µM acetaldehyde or acetate, or into the same medium without these compounds (control), at an initial OD₆₀₀ of 0.0005. Cultures were incubated at 22°C for 4 days. To compensate for acetaldehyde evaporation during cultivation, an additional 50 µM acetaldehyde or acetate was added every 24 h, and 100 µL of culture supernatant was sampled at the same time points. Growth curves of the mutant and wild-type strains were generated by determining viable cell densities via colony counts after spreading serial dilutions on R2A agar plates. Each condition (acetaldehyde, acetate, and control) was tested with three biological replicates.

The colonization ability of the strain DW067ΔALDH mutant and wild-type strain was evaluated under mono-inoculation (six replicates) and within a six-member SynCom (three replicates), following exactly the same procedure as Ishizawa et al. (2024a). Colonization densities of the wild-type and mutant strains were compared using Welch’s *t*-test in R.

### Determination of acetaldehyde content

Sterile duckweed was cultivated for 10 days using the same method as that used for the transcriptome analysis, with four biological replicates. Acetaldehyde concentrations in the surrounding water were measured using an Acetaldehyde Assay Kit (Sigma-Aldrich).

## Data availability

All sequence reads generated during this study were deposited in the DDBJ/EMBL/GenBank database under BioProject accession number PRJDB12961. All datasets and analytical codes are available in the Zenodo repository, doi: 10.5281/zenodo.18382841.

## ACKNOWLEDGEMENTS

This work was funded by the Japan Society for the Promotion of Science KAKENHI grant numbers JP20J00210, JP23K14268, and JP22H04925 (PAGS), JST PRESTO grant number JPMJPR25N1, Nakatsuji Foresight Foundation Research Grant, and the Technology Research Partnership for Sustainable Development (SATREPS) (JPMJSA2004) in collaboration between the Japan Science and Technology Agency (JST) and the Japan International Cooperation Agency (JICA).

