## Supplementary Figures for "Competition-induced metabolic switching governs the interaction network in plant microbiome assembly"

*Corresponding author

Hidehiro Ishizawa

Address: C621 Shosha 2167, Himeji, Hyogo, 671-2280 Japan,

Phone: (+81)79-267-4969

Mail to:

**Contents**


- **Supplementary Figure S1**
  Non-metabolic response of the six SynCom members
- **Supplementary Figure S2**
  Exometabolomic analysis of Novosphingobium olei DW067 and its ALDH gene mutant.
- **Supplementary Table S1**
  Analytical conditions for GC-MS analysis
- **Supplementary Table S2**
  Parameters used for metabolome data processing in MS-DIAL
- **Supplementary Table S3**
  Strains, plasmids, and primers used for constructing ALDH knockout

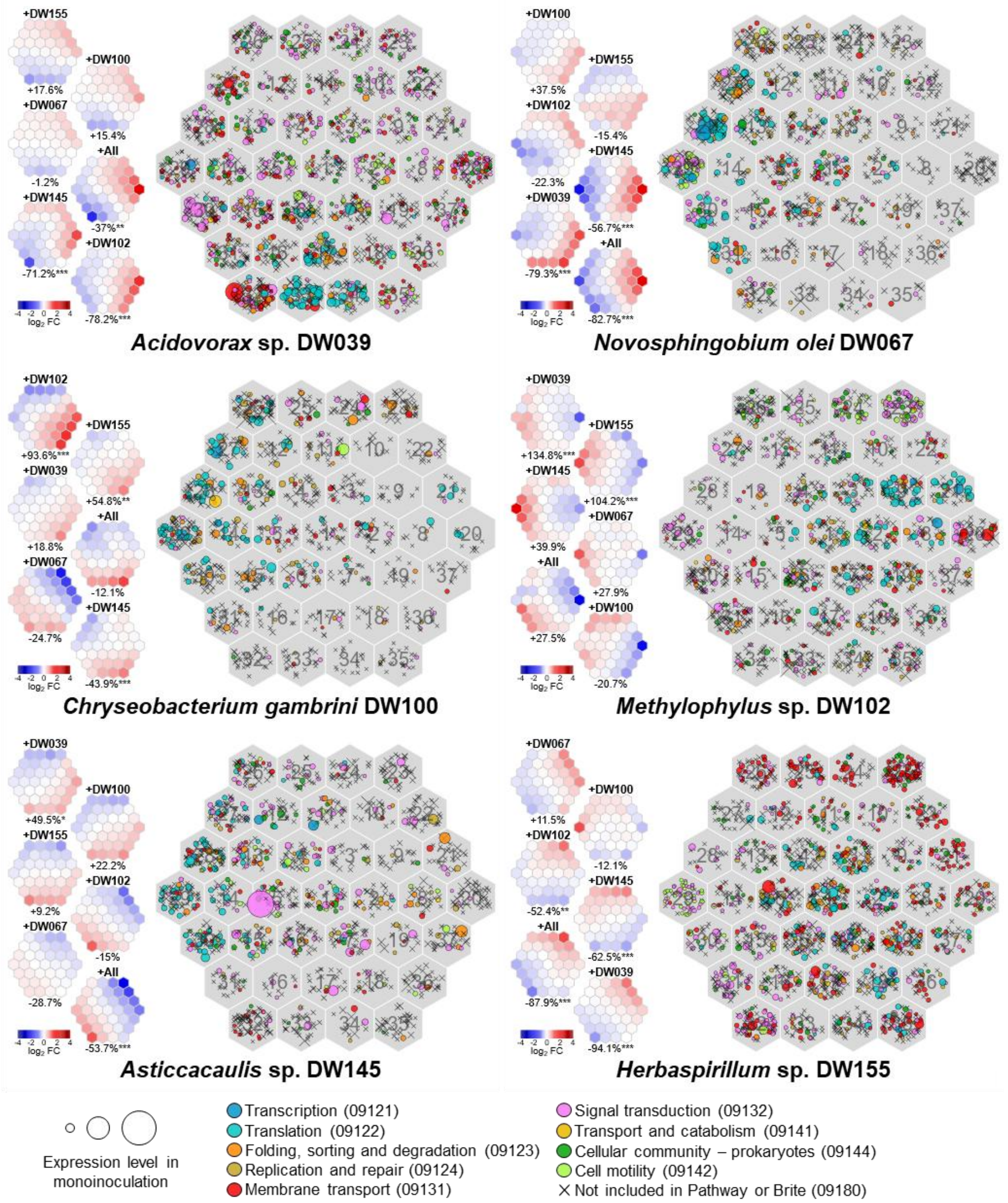


**Fig. S1** Non-metabolic responses of the six SynCom members. Each SynCom member was inoculated onto sterile duckweed under three conditions: mono-inoculation, pairwise co-inoculation with each of the five other SynCom members, and co-inoculation with all five strains, after which total mRNA was sequenced. The hexagonal maps show the results of self-organizing map (SOM) clustering based on gene expression patterns. Each hexagon contains genes assigned to the same SOM cluster; symbol colors indicate KEGG functional categories, and symbol sizes represent expression levels under mono-inoculation. The heatmaps on the left show the mean expression level (fold change relative to mono-inoculation) of genes in each SOM cluster. The percent change in colonization density of the focal strain relative to mono-inoculation is shown below each heatmap as a measure of interspecies interaction from co-inoculated strains.

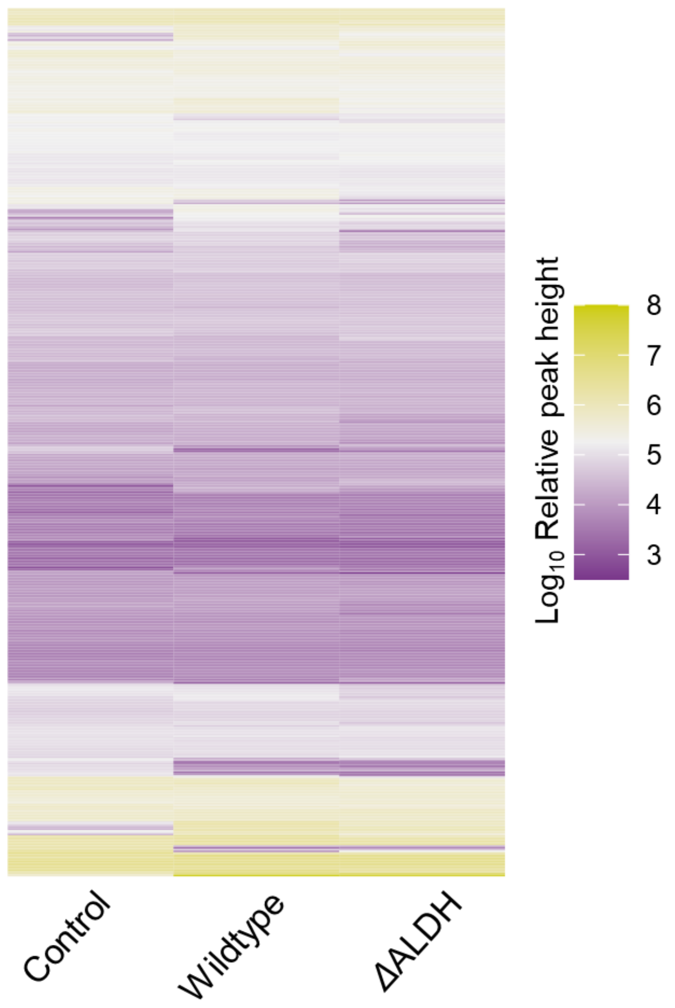


**Fig. S2** Exometabolomic analysis of *Novosphingobium olei* DW067 and its ALDH gene mutant. The wild-type and mutant strains were each grown in extracted duckweed exudates, and the resulting culture residues were analyzed by untargeted GC–MS after trimethylsilyl derivatization. Relative intensities of 505 detected metabolite peaks are shown.

**Table S1** Analytical conditions for GC-MS analysis

| Items | Conditions |
| --- | --- |
| *Gas chromatography* | |
| Injection temperature | 250°C |
| Column temperature | 80°C (2 min) - 320°C (20 min) |
| Injection mode | Splitless |
| Carrier gas | He (constant linear velocity, 1 mL min$^{-1}$) |
| Linear velocity | 39 cm sec$^{-1}$ |
| Split ratio | -1.0 |
| Injection volume | 1 µL |
| *Mass spectrometry* | |
| Ion source temperature | 200°C |
| Interface temperature | 250°C |
| Solvent delay time | 2.5 min |
| Detector voltage | 0.2 kV (Relative to the Tuning Result) |
| Start time | 4.50 min |
| Finish time | 25.00 min |
| Event time | 0.2 sec |
| Scan speed | 5000 µ sec$^{-1}$ |
| Scan range | *m*/*z* 30-500 |

**Table S2** Parameters used for metabolome data processing in MS-DIAL

| **Parameter** | **Value** |
|---|---|
| *Data collection* | |
| Mass scan range | 30 – 500 Da |
| *Peak detection* | |
| Minimum peak height | 1000 amplitude |
| Smoothing method | Linear weighted moving average |
| Smoothing level | 4 scan |
| Average peak width | 20 |
| Mass slice width | 0.5 |
| Mass accuracy | 0.5 |
| *MS1Dec* | |
| Sigma window value | 0.5 |
| EI spectra cut off | 10 amplitude |
| *Identification* | |
| Retention index tolerance | 30 |
| *m*/*z* tolerance | 0.5 Da |
| EI similarity cut off | 70% |
| Identification score cut off | 70% |
| Use retention information for scoring | True |
| Use retention information for filtering | True |
| Use quant masses defined in MSP format file | True |
| *Alignment* | |
| Retention index tolerance | 30 |
| EI similarity tolerance | 70 |
| Retention time tolerance | 0.1 |
| EI similarity factor | 0.5 |
| Identification after alignment | False |
| Gap filling by compulsion | True |
| Basepeak *m*/*z* selected as the representative Quant mass | False |

**Table S3** Strains, plasmids, and primers used for constructing ALDH knockout

| Items | Relevant characteristics or sequences (5' – 3') | Source |
|---|---|---|
| *Strains* | | |
| *Novosphingobium olei*. DW067 | Duckweed-associated bacteria, Streptomycin resistant | Ishizawa et al. (2020a) |
| *Escherichia coli* DH5α | Cloning host | Takara Bio |
| *Escherichia coli* HST08 | Cloning host | Takara Bio |
| *Plasmids* | | |
| pK18mobsacB | Suicide vector, Kanamycin resistant, *lacZ* fragment, *sacB* | Schäfer et al. (1994) |
| pRK2013 | Helper plasmid, Kanamycin resistant, *tra* | Biomedal |
| *Primers* | | |
| pk18F | CTGAATGGCGAATGGCGATAAGC | this study |
| pk18R | ACCGAGCTCGAATTCGTAATCATG | this study |
| DW067aldh_ins1F | ATTACGAATTCGAGCTCGGTAAGCTGTACCACGCGCTC | this study |
| DW067aldh_ins1R | CCGAACGAGAGTACTTGTCGAGGCCTTGAGG | this study |
| DW067aldh_ins2F | CGACAAGTACTCTCGTTCGGGTTCTTGTGG | this study |
| DW067aldh_ins2R | TATCGCCATTCGCCATTCAGTGCATCATGGCCTCGATCC | this study |
